# OPTIMIZING THERAPEUTIC TARGETS FOR BREAST CANCER USING BOOLEAN NETWORK MODELS

**DOI:** 10.1101/2023.05.10.540187

**Authors:** Domenico Sgariglia, Flavia Requel Gonçalves Carneiro, Luis Alfredo Vidal de Carvalho, Carlos Eduardo Pedreira, Nicolas Carels, Fabricio Alves Barbosa da Silva

## Abstract

Studying gene regulatory networks associated with cancer provides valuable insights for therapeutic purposes, given that cancer is fundamentally a genetic disease. However, as the number of genes in the system increases, the complexity arising from the interconnections between network components grows exponentially. In this study, using Boolean logic to adjust the existing relationships between network components has facilitated simplifying the modeling process, enabling the generation of attractors that represent cell phenotypes based on breast cancer RNA-seq data. A key therapeutic objective is to guide cells, through targeted interventions, to transition from the current cancer attractor to a physiologically distinct attractor unrelated to cancer. To achieve this, we developed a computational method that identifies network nodes whose inhibition can facilitate the desired transition from one tumor attractor to another associated with apoptosis, leveraging transcriptomic data from cell lines. To validate the model, we utilized previously published in vitro experiments where the downregulation of specific proteins resulted in cell growth arrest and death of a breast cancer cell line. The method proposed in this manuscript combines diverse data sources, conducts structural network analysis, and incorporates relevant biological knowledge on apoptosis in cancer cells. This comprehensive approach aims to identify potential targets of significance for personalized medicine.

## INTRODUCTION

Cancer is a disease characterized primarily by uncontrolled cellular proliferation. This dysregulation disrupts normal cellular homeostasis, leading to the emergence of distinctive traits known as “hallmarks of cancer,” which are common across different tumor types [1]. Carcinomas, a type of epithelial cell tumor, account for approximately 85% of all cancers and can affect various tissues in the human body. When these tumors occur in glandular tissue, they are specifically referred to as adenocarcinomas. Breast cancer falls into the category of adenocarcinomas. It is the most prevalent neoplastic condition affecting women, with a global incidence of approximately 2,261,419 new cases and 684,996 deaths in 2020 [2].

In addition, treating this pathology gives rise to harmful adverse effects in patients. For instance, a study identified 38 distinct negative symptoms categorized into five groups that resulted from chemotherapy administration [3]. Therefore, developing new intervention strategies that can enhance therapies and minimize their unwanted side effects is crucial. We propose using a Boolean modeling approach for breast cancer to address this need. Cancer is a genetic disease with multifaceted ramifications [4]. Cancer cells’ DNA undergoes numerous alterations due to the oncogenic process, including single-base pair mutations, indels, and epigenetic modifications.

Cancer occurrence leads to network modifications, where the pathways involved are frequently intertwined to generate processes characteristic of tumor dynamics and progression [5-7]. Epigenetic changes and alterations in gene regulatory networks [8] provide an opportunity for modeling cancer attractors [9]. This study builds upon a previous analysis [10] of attractors identified within a gene regulatory network based on breast cancer data. Specifically, by incorporating a novel set of genes associated with apoptosis into the Boolean network, we identified new attractors resulting from target inactivation. This modeling enabled our gene model to transition towards a cell death phenotype, as observed in corresponding *in vitro* experiments [11].

This paper presents an algorithm that optimizes the selection of network elements capable of inducing trajectories between attractors in the epigenetic landscape. Additionally, we have introduced an indicator that quantifies the network’s response when inducing a trajectory from a malignant state to an apoptosis state through direct intervention on its vertices. We incorporated a set of genes representing the apoptosis process into the gene regulatory network associated with breast malignancy to achieve this. By manipulating the activation or inhibition state of each gene in this group, we assessed the effectiveness of network perturbations in transitioning the phenotype from malignancy to apoptosis. We calibrated the network based on (i) the typical gene expression level observed in the malignant attractor and (ii) the genes to be inhibited for inducing apoptosis in a malignant cell line, as determined from *in vitro* experiments. To validate the system dynamics, we compared the results with the *in vitro* experiment [11], where five genes were silenced to induce the death of a breast cancer cell line. This experimental data was used to evaluate the network’s behavior upon the permanent silencing of specific targets.

We confirmed that the network structure derived from the interactome could drive the malignant attractor toward apoptosis by selectively silencing the same network vertices as those targeted in the *in vitro* experiment. The capacity of our model to replicate the conditions conducive to malignant cell death observed *in vitro* enabled us to optimize the selection of targets for transitioning the system dynamics from the malignant attractor to the apoptosis state. This optimization process involved utilizing specific analysis techniques to examine the network structure, enabling us to identify the vertices whose inhibition could mimic and enhance the outcomes achieved in the *in vitro* experiment.

## MATERIALS AND METHODS

The various stages involved in conducting this research are briefly outlined in Figure 1.

**Fig. 1:**
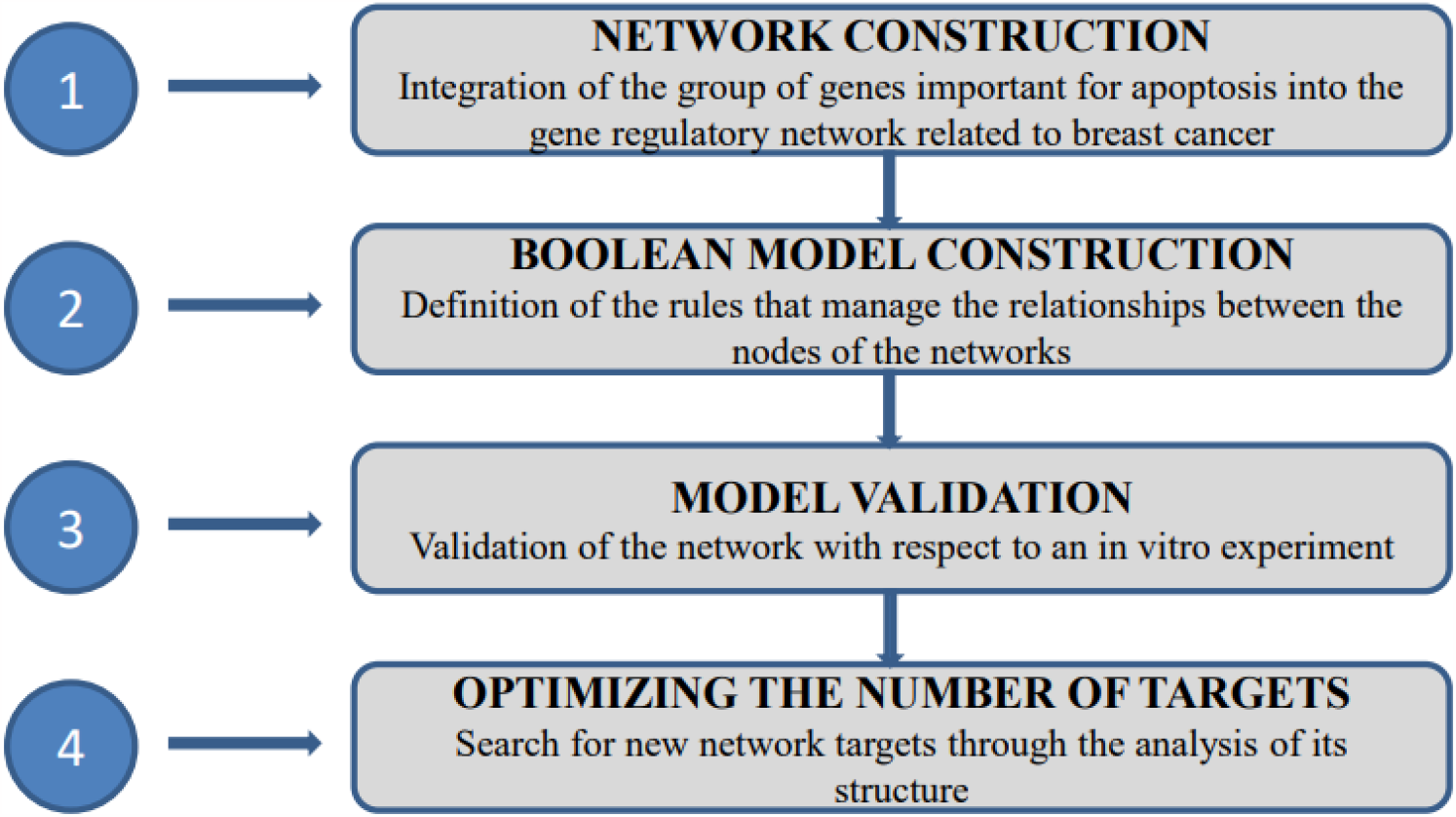
Steps for Boolean network construction and dynamic simulation.

**Fig. 2:**
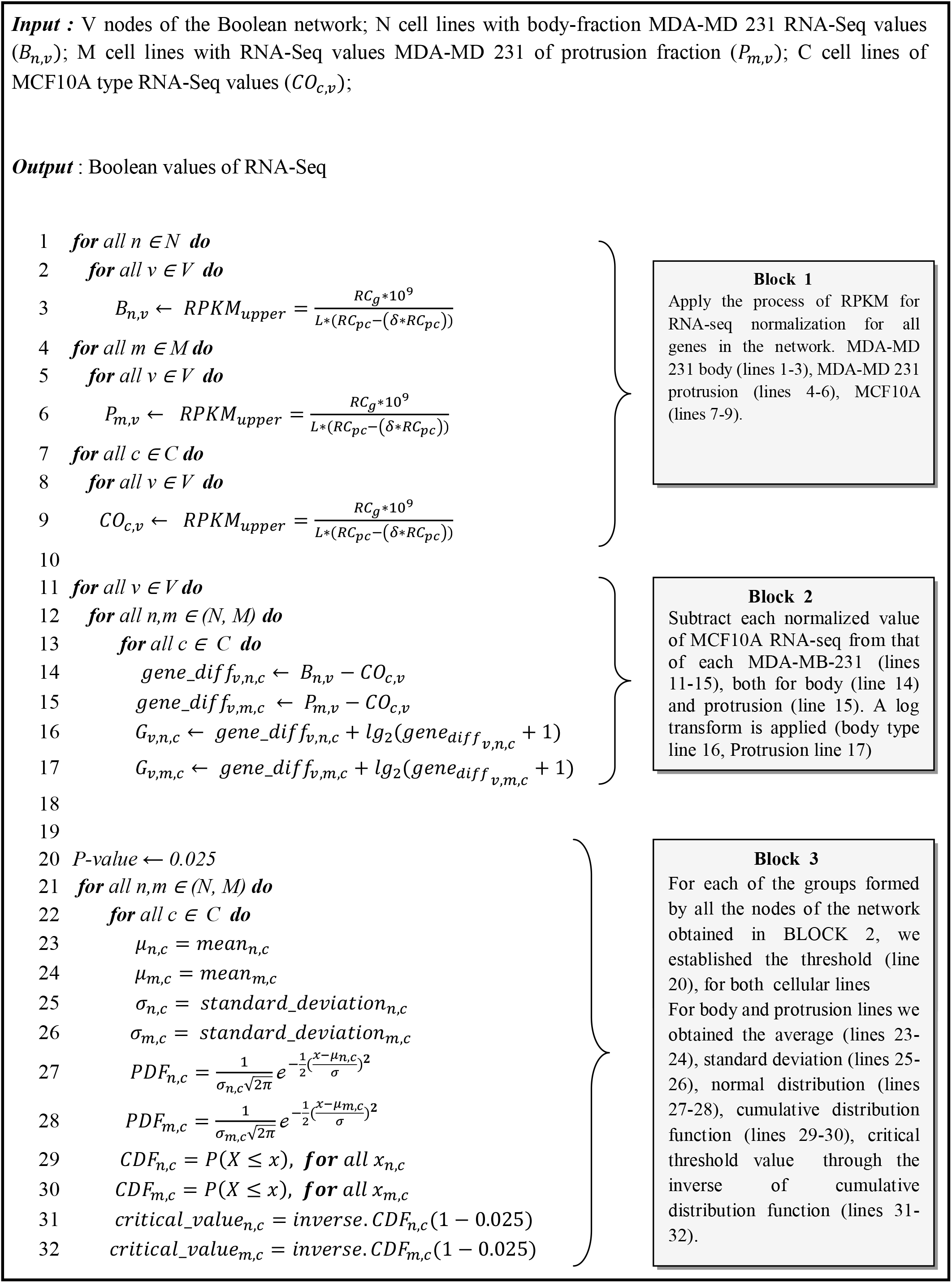
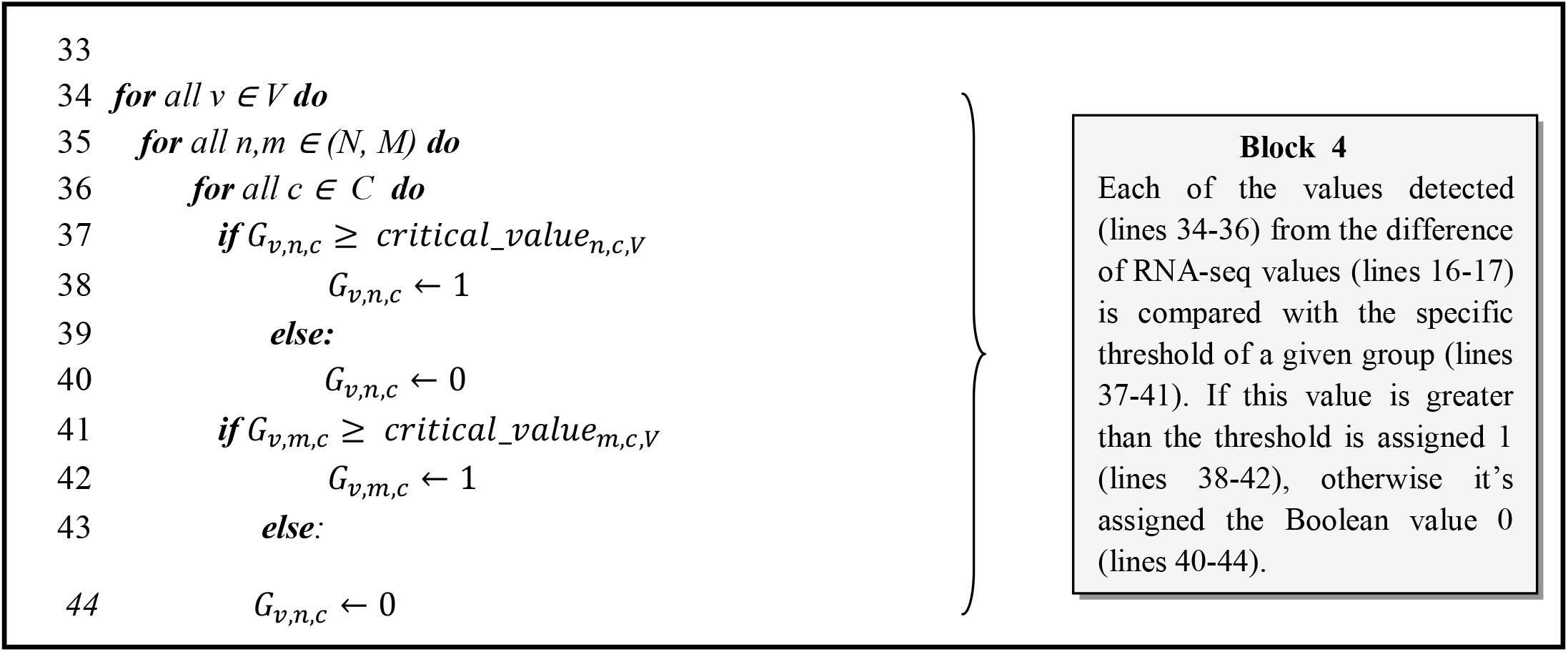
Procedure used for binarization of RNA-seq values. BLOCK 1: normalization of RNA-seq values. BLOCK 2: subtraction of each normalized value of MCF10A RNA-seq from each value of MDA-MB-231data and log transformation application. BLOCK 3: determination of the threshold value for which to attribute a specific Boolean value. BLOCK 4: attribution of a Boolean value based on the critical value.

### 1 NETWORK CONSTRUCTION

We initiated our study using a gene regulation network established in a previous publication [10]. Breast cancer RNA-seq data guided the selection of network nodes, and the chosen genes were linked to their respective hallmarks using the MSigDB repository. Notably, the two crucial hallmarks of cancer, namely “UNLIMITED REPLICATIVE POTENTIAL” and “EVASION OF CELL DEATH,” were well-represented in the dataset. We introduced an additional set of 28 genes to supplement the initial network consisting of 103 genes [10] to enlarge the network. Twenty-five incorporated nodes were for apoptosis-associated genes, exerting either inducing or inhibitory effects on this cellular process. Alongside these 25 novel vertices, two existing vertices from the previous study assume a crucial role in cellular apoptosis as constituents of the apoptotic cascade. Collectively, we refer to these 27 genes as apoptosis-related genes. This network enlargement was imperative to facilitate the modeling of the transition from the malignant state to the apoptosis one, which was induced by network perturbations through targeted inactivation of specific vertices. Protein-protein interactions were acquired from the IntAct interactome (IntAct database, version updated in December 2017) to establish the network connections between genes. The specific file used for obtaining the interactions was retrieved from the FTP link ftp://ftp.ebi.ac.uk/pub/databases/intact/current/psimitab/intact-micluster.txt, accessed on January 11, 2018. The directionality of the connections and their regulatory nature (activation or inhibition) were determined by consulting the Metacore database [12]. To verify the network’s validity in replicating a tested biological scenario *in vitro*, our system was configured to reproduce the outcomes documented by Tilli et al. [11]. Their experimental study demonstrated cell death in a cancer cell line (MDA-MB-231) by inhibiting five genes using RNA interference. To achieve this objective, we retrieved the RNA-seq data of two distinct cell lines, namely MCF10A and MDA-MB-231, from the Gene Expression Omnibus (GEO) repository available at https://www.ncbi.nlm.nih.gov/gds/. MDA-MB-231 is a malignant cell line derived from triple-negative breast cancer, while MCF10A served as the non-tumoral control in this study. For MCF10A, we obtained the following RNA-seq datasets: SRR2149928, SRR2149929, SRR2149930, SRR2870783, and SRR2872995. Regarding MDA-MB-231, we acquired the RNA-seq datasets: ERR493677, ERR493680 (corresponding to the *body* portion), and ERR493678, ERR493679 (corresponding to the *protrusion* portion) of the cells.

Based on information obtained from the GEO repository, the cells were cultured on a polycarbonate transwell filter with 3-micrometer pores, allowing the formation of protrusions through the pores for 2 hours. Subsequently, the cells underwent a washing step, and both sides of the filter were lysed to extract RNA for further analysis. Through this protocol, the cells were fractionated into two distinct fractions: *protrusion* and *body* types. In the subsequent analysis, we employed a binary approach to categorize the up-regulated genes observed in each MDA-MB-231 RNA-seq dataset (both *body* and *protrusion*) compared to every MCF10A RNA-seq dataset. This approach introduced variability into the experiment and facilitated the assessment of system robustness.

To ensure the convergence of our network model with scale-free networks, which is characteristic of cell signaling pathways [19], we examined the degree distribution of the network vertices. To evaluate the network’s structure, we compared its degree distribution with that of random graphs [27], Watts and Strogatz small-world networks [28], and scale-free networks [19], all generated with the same number of vertices. To facilitate this comparison, we utilized the complementary cumulative distribution function (CCDF) as defined by equation 1. The CCDF provides the probability (F) of a vertex having a connectivity degree equal to or greater than a specified value. By analyzing these distributions, we could determine the degree of alignment between our network and these reference models.

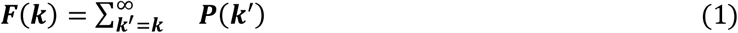

### 2 BOOLEAN MODEL CONSTRUCTION

After constructing our model’s directed graph, we established the conditions necessary for its dynamic simulation by defining the transfer functions that govern the system’s evolution at discrete time intervals. The objective was to guide the dynamic behavior of the network elements in a manner that faithfully replicated the observed conditions from the *in vitro* experiment conducted by Tilli et al. [11]. By incorporating these specific conditions, we aimed to ensure the accurate representation of the experimental findings within our model’s dynamic framework. In systems biology, accurately deducing the interaction rules of a network poses a significant challenge. To address this complexity, we employed Boolean nested canalizing functions [13], where the function is influenced by the specific order in which variables are organized. A Boolean function is considered canalizing if a single input can solely determine the output. In cases where this input does not play the canalizing role, the other inputs are deemed responsible for fulfilling this function. By adopting the hierarchical structure of transfer functions, achieved through nested canalizing functions, we aimed to capture the behavior of biological systems more effectively [14,15]. Furthermore, many network nodes exhibited a substantial number of inputs, emphasizing the need for a robust modeling approach. In this scenario, using nested canalizing functions offers increased system stability [16], which is crucial in managing the inherent noise observed in biological systems.

The utilization of nested canalizing functions and the need to align the model with biological facts enabled us to manually establish the transfer rules for each gene [16,17]. Opportunely, Harris et al. [18] demonstrated that a significant portion of the gene updating rules fell under canalizing functions. Considering these considerations, we determined the number of inputs for the Boolean functions, as defined in Equation 2.

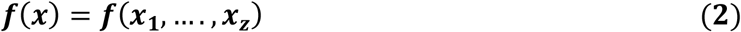

Considering a set of Boolean variables *x* = {*x*_1_, *x*_2_,…,*x*_z_}, the input was defined as essential (see equation 3) if the condition of equation 3 was satisfied.

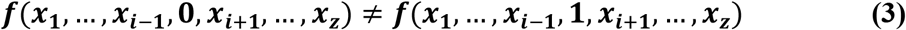

The essential inputs were defined as *canalizing* if there were values *a,b* ∈ {0,1}that satisfied equation 4 for all remaining combinations of variables x-{*x*_*i*_}where *x* is a canalizing input value, and *x*_i_ is a canalized value.

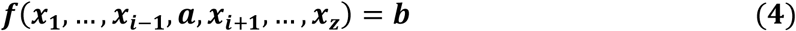

A function with ***z*** essential input is defined as nested if it is z-times canalized with *x*_1_,…,*x*_*z*_ canalizing inputs and ***a***_**1**_,…,***a***_***z***_ canalizing values, to which correspond the canalized value ***b***_**1**,…,_***b***_***z***._

Nested canalizing functions are an extension of the canalizing function formalism [13], in which the order of the inputs is considered to assign the canalizing role. In addition, canalizing functions can be nested if it is possible to set them with **z** inputs and **z -1** Boolean operators AND (**∧)** or OR (**⋁)** with a priority proceeding from left to right. Thus, defining (*∈{∧ ⋁}), we have equation 5.

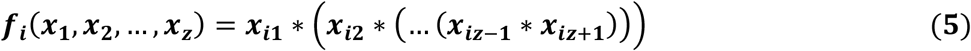

In the implementation phase of this report, each input of the function was coupled to a single logical operator, which can be an **∧ and** or an **⋁or**. In light of these rules, the transfer functions of the network have been implemented according to Equation 6

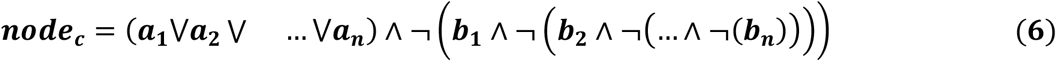

where ***node***_***c***_ is the *n*^th^ node of the network, ***a***_***i***_, ***i* =[1**,***n*], *n*** represents the number of nodes with activation function on ***node***_***c***_, and ***b***_j_, ***j* =[1**,***m*] *m*** is the number of nodes with inhibition function on . To create suitable conditions for the network to reproduce the results obtained in the *in vitro* experiment [11], we applied some changes to the general scheme of the nested canalizing functions illustrated above in the nodes representing the *TP53, HIF1A, RELA, NFKB1, HDAC1, STAT3, BCL2, CASP3*, and *BRCA1* genes as shown in equations 7 to 11 where some input variables with activation or inhibition roles on *node*_*c*_ ceased to be independent of the other elements of the function and assume a cumulative role for the final result, which the other nodes cannot replace.

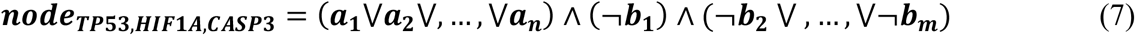

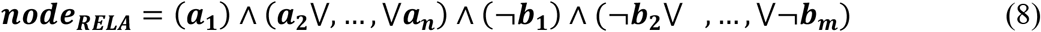

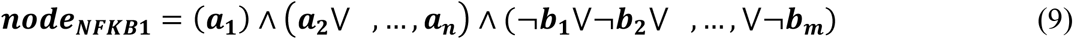

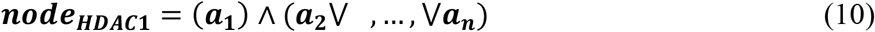

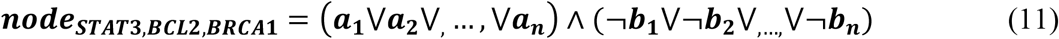

After defining the constituent elements of the gene regulation network and analyzing its structure, the subsequent task was determining the appropriate mathematical formalism for the system’s dynamic analysis. We opted to employ a directed graph model based on Boolean logic. Boolean network modeling represents one of the simplest methods for dynamic modeling while offering the advantage of reliably providing insights into system dynamics.

In this context, we considered a Boolean variable, denoted as ***B***, which takes on the value of True (1) or False (0) depending on whether a particular gene is up-regulated or not in the RNA-seq data of the MDA-MB-231 (malignant) cell line compared to the MCF10A (control) cell line. Consequently, for the ***n*** vertices within our network, we can express this relationship using equation 12.

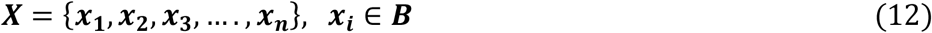

When time is represented as a discrete scalar value, the states of the network can be depicted as a vector with its components being the vertices of the network (equation 13).

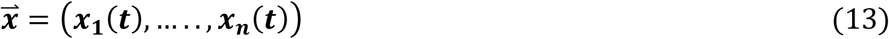

The trajectories of the system within the state space are then contingent upon the Boolean functions associated with the n vertices of the network (equation 14).

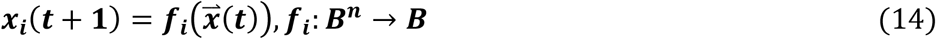

In this report, we used a synchronous update mode for the network vertices, wherein all vertices are updated simultaneously. While an asynchronous update mode may align better with biological realism, the choice of update mode is not crucial given the computational and conceptual advantages of synchronous updates and the enhanced system stability achieved through the utilization of nested canalized transfer functions [21]. In the synchronous update mode, the system’s progress occurs in consecutive temporal states (equation 15).

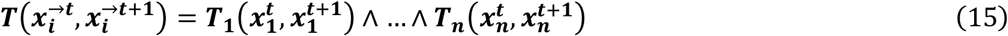

The goal of Boolean modeling is to identify the attractors expressed by the dynamics of the system. Attractors are stable gene activity patterns that represent the long-term behavior of the Boolean network and are interpreted as a specific cellular phenotype.

Attractors in a Boolean system can be subdivided into different classes. Examples are fixed-point attractors, characterized by a single state of the system (i.e., the Boolean configuration of network nodes) that persists indefinitely, and cyclic attractors, characterized by a sequence of states that repeat periodically. Each attractor is matched with a specific basin of attraction, composed of all the system states for which it represents the stable state at the end of their dynamic evolution.

### 3 MODEL VALIDATION

Initially, we compared MDA-MB-231 RNA-seq samples (two from the *body* and two from the *protrusion*) and each corresponding MCF10A sample. To achieve this, we employed the Reads per kilobase of transcript per Million reads mapped (RPKM) normalization process, as outlined in Pires et al. [20], to normalize the read counts of the twenty paired RNA-seq samples. Subsequently, we subtracted each normalized value of the MCF10A RNA-seq sample from the corresponding MDA-MB-231 data. For positive values (indicating up-regulated genes in the malignant state), we applied a logarithmic transformation based on equation 12, utilizing the pipeline described by Pires et al. [20].

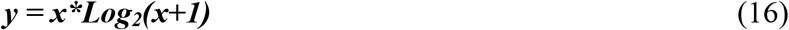

The procedure involved in this pipeline consists of classifying genes as up-regulated based on whether the logarithmic transformation of their differential expression surpasses a critical threshold. To determine this critical value, a Python script was employed to fit a Gaussian curve with a 95% confidence level to the data for a p-value of 0.025 (for more details, refer to Pires et al., 2021). In each of the twenty comparisons between MBA-MD-231 and MCF10A, genes that were identified as up-regulated through this process were assigned a value of “1”, while the remaining genes were assigned a value of “0” (**Supplementary Table S1**).

Using the BooleanNet library [22], we analyzed the binary values of each MDA-MB-231 RNA-seq normalized data in conjunction with the corresponding genes from every MCF10A sample within our network (**Supplementary Table S2**). Our objective was to identify the attractors generated through the dynamic evolution of the network. The presence of the 27 apoptosis-related genes in the attractors of this initial configuration served as a reference point for evaluating the impact of subsequent network modifications (Figure 3).

**Fig. 3:**
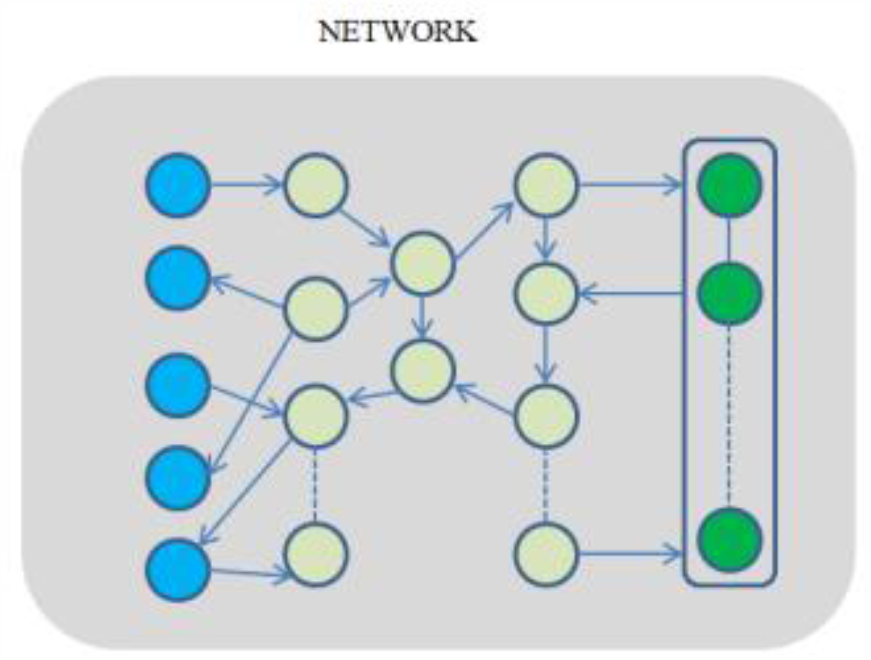
Structure of the gene regulation network under study. The blue color indicates the five target genes, *CSNK2B, HSP90AB1, TK1, VIM, YWHAB*, which were inhibited in the *in vitro* experiment of Tilli et al. [11]. The dark green nodes represent the apoptosis-related genes. The nodes in light green exemplify the rest of the network genes. There is no vertex inhibition in this network representation.

Subsequently, we conducted simulations to replicate the conditions of the *in vitro* experiment performed by Tilli et al. [11], where the MDA-MB-231 cancer cell line’s death was induced through transient inhibition of *TK1, VIM, YWHAB, CSNK2B*, and *HSP90AB1* genes using siRNA interference. For the sake of clarity, we will, below, refer to these five targets as *bench targets*. To emulate this *in vitro* experiment, we permanently inhibited the vertices corresponding to these bench targets in the dynamic evolution of the network, as shown in Figure 4.

**Fig. 4:**
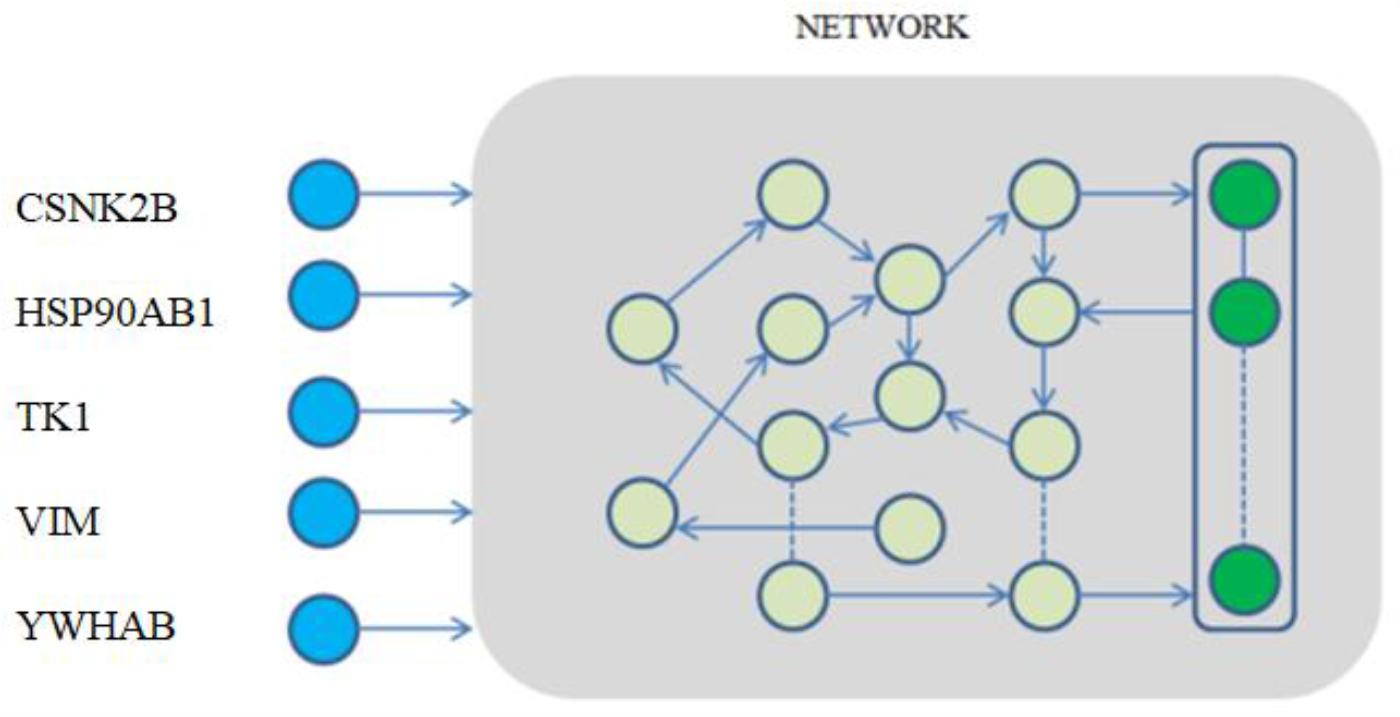
The five bench targets (blue nodes) were set to zero (inhibition) for network dynamics simulation.

Figure 4 compares the activation or inhibition of the 27 apoptotic genes and their involvement in the attractors after inhibiting the five bench targets in the original network depicted in Figure 3. This comparison allowed us to evaluate the functional agreement between our model and the *in vitro* experiment conducted by Tilli et al. [11]. Furthermore, TP53 was permanently inhibited throughout the network simulation since mutations render it ineffective as a tumor suppressor in MDA-MB-231 [41].

### 4 OPTIMIZING THE NUMBER OF TARGETS

Based on the *in vitro* induction of cell death in MDA-MB-231 through the silencing of *CSNK2B, HSP90AB1, TK1, VIM*, and *YWHAB* [11], and considering our understanding of the key 27 apoptosis-related genes, we present a methodology to identify genes capable of driving cancer cells towards programmed cell death.

To maximize the presence of the 27 apoptosis-related genes within the apoptosis attractor, with genes promoting apoptosis being activated and genes inhibiting it being deactivated, our objective was to identify the most effective target genes within the network structure. To achieve this, we examined the network’s modularity based on the connectivity of its vertices. The Clauset-Newman-Moore greedy modularity maximization algorithm [23] was used to identify the modular structure, considering the network as undirected. Modularity was calculated using equation 17, where *c* represents communities, *L*_*c*_ denotes the number of links within a community, is the resolution parameter, and is the sum of degrees within the community.

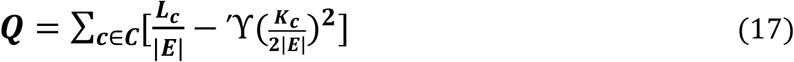

In the initial stage of the algorithm, each node is assigned to its own cluster, forming a partition. The algorithm then proceeds iteratively, merging pairs of clusters to increase modularity. The initial modularity value is negative, representing a singleton cluster, and gradually increases until reaching a positive peak, corresponding to the optimal solution found by the algorithm. Eventually, the modularity value returns to zero when all nodes are in the same community. In a backward process, the algorithm identifies the partition corresponding to the peak value. The implementation of this algorithm utilized the NetworkX Python library [24]. After detecting the communities in the network, we examined whether the 27 apoptosis-related genes were grouped or dispersed among these identified modules, with the possibility of forming a single community by the algorithm’s criteria. This approach has previously been implemented in [25], where a modularized network was used to map drug targets for cancer and identify modules that were the focus of therapeutic action. Due to the observed clustering of apoptosis-related genes in the network structure, we searched nodes that could be bridges among these apoptosis-related clusters and the five bench targets. Identifying these bridge nodes could potentially shorten the path for target genes to replicate the outcomes of the *in vitro* experiment. To accomplish this, we employed the Dijkstra algorithm using the NetworkX library in Python [24] to find the shortest path between each of the five bench targets (*CSNK2B, HSP90AB1, TK1, VIM, YWHAB*) and every apoptosis-related gene.

A similar approach was adopted by George et al. [26], wherein intermediate genes along the shortest path were identified as potential therapeutic targets. These genes were then ranked based on the number of shortest paths in which they were involved.

Utilizing the knowledge gained from the *in vitro* experiment, which demonstrated that inhibiting the five bench targets resulted in cell death of MDA-MB-231, we sought alternative vertices that, when inhibited, would activate the maximum number of genes within the apoptosis group. To accomplish this, we aimed to identify the smallest set of vertices that shared the common property of being involved in at least one shortest path between each of the five genes (*CSNK2B, HSP90AB1, TK1, VIM, YWHAB*) as starting nodes and any apoptosis-related gene as the final node (Figure 5).

**Fig. 5:**
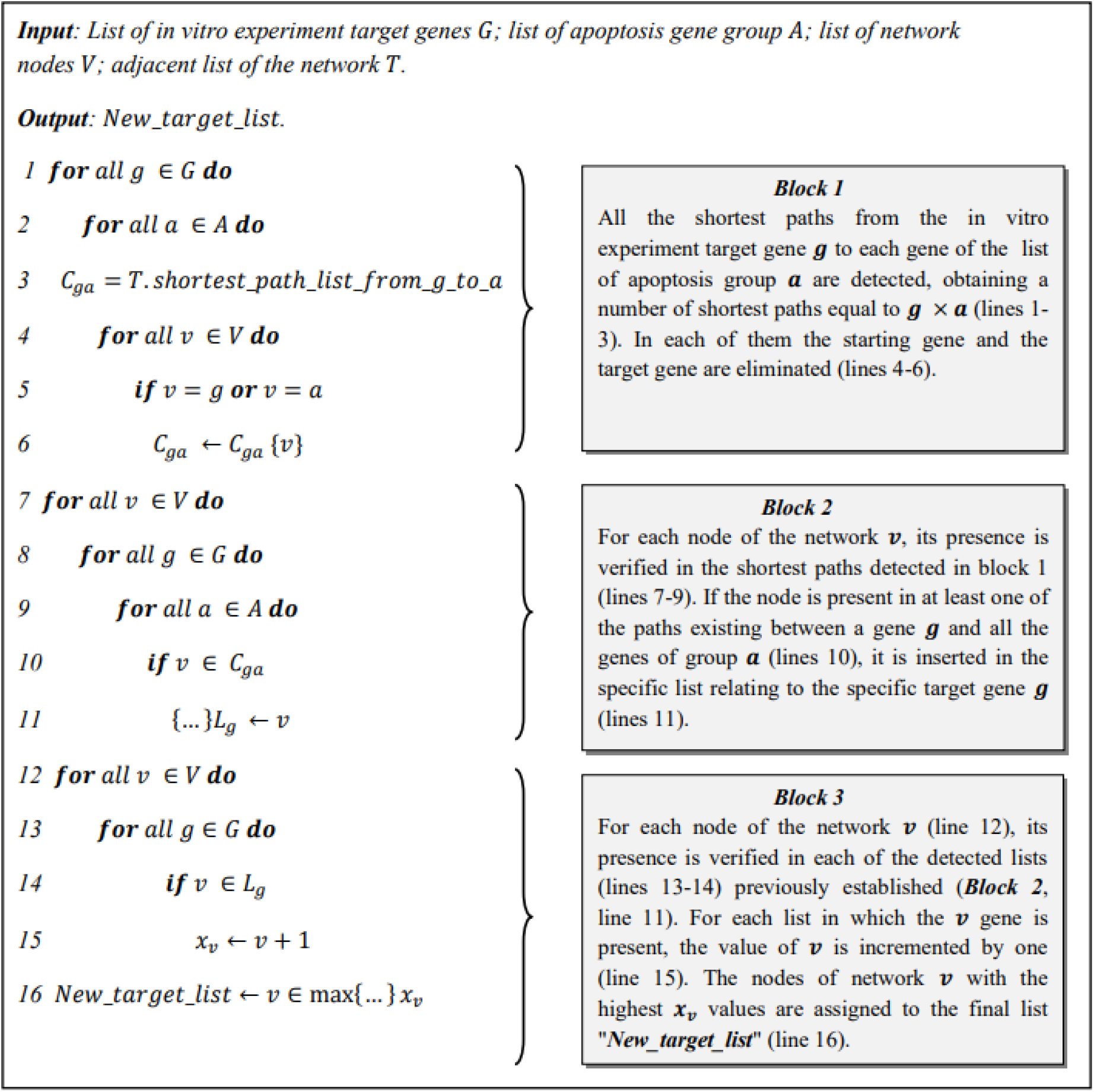
Pseudocode of target vertex determination in shortest paths. Lines 1-6 (block 1): Detection of the shortest paths via Python networkX library between the five bench targets and the apoptosis-related genes. Lines 7-11 (block 2): Creation of a list for each bench target gene containing the nodes detected by the algorithm on every shortest path. Line 12-17 (block 3): Insertion in the list of new targets of the genes present in the largest number of lists of the previous step. The asymptotic complexity of the algorithm is *O* (max{*C*_1,2_(*G*\* *A*\**V*), *C*_3_(*V*\**G*)}).

The methodology outlined in Figure 5 enabled us to identify novel target genes that can potentially influence the configuration of the apoptosis-related group. By “configuration,” we refer to both the count of activated apoptosis-related genes that can induce apoptosis and the consistency of this count across the RNA-seq comparisons conducted in this study.

To evaluate the impact of deactivating the newly identified target vertices using the shortest path strategy on the configuration of the apoptosis-related group, we examined the activation or inhibition status of the apoptosis-related genes across the twenty analyzed comparisons (ten for the *body* and ten for the *protrusion*, representing the two fractions of MDA-MB-231). This approach enabled us to compare the effects of inhibiting the new targets within the shortest paths to those obtained by inhibiting the original five bench targets.

## RESULTS

### 1 GENE REGULATORY NETWORK

The breast cancer regulatory network utilized in this study consists of the genes employed in a previous report for the computation of Boolean attractors [10], which were further expanded by the inclusion of 25 additional genes (Fig. 6). These 25 genes, along with the two genes already present in the initial network, play a key role in the cellular apoptosis process.

**Fig. 6:**
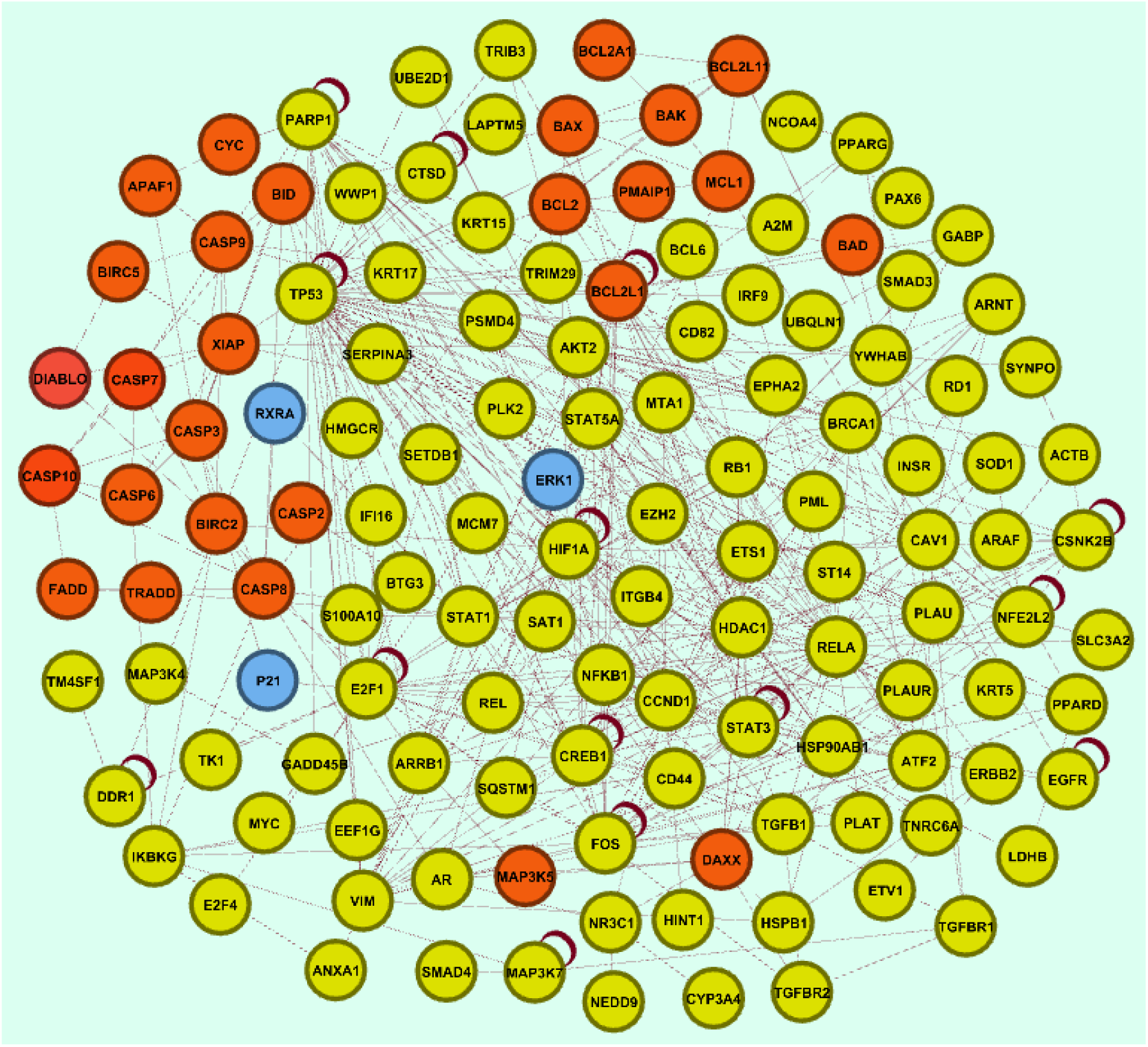
Breast cancer gene regulatory network used in this report. Twenty-five red nodes and the three blue nodes, not part of the apoptosis group, are the new vertices added to the network used in the previous report (yellow nodes). In the group of twenty-seven red genes related to apoptosis, two (*BAX* and *BCL2L1*) were already present in our earlier work network.

The network depicted in Figure 6 consists of **131** nodes and **494** edges. The system’s dynamics are governed by Boolean transfer functions, where each node can act as either an activator or an inhibitor on the other nodes to which it is connected (**Supplementary Table S3**). The nature of these interactions was deduced using a dedicated software [12], which also facilitated the integration of the genes from the apoptosis-related group into the pre-existing network.

### 2 STRUCTURAL ANALYSIS OF THE NETWORK

We examined the structural features of the network by comparing it to well-known canonical network types. This comparison involved analyzing the degree distribution and comparing it to three distinct network types commonly described in the literature: The Erdos-Renyi network [27], the Watts-Strogatz network [28], and the Barabasi-Albert network [19].

By analyzing the plot presented in Figure 7, which depicts the complementary cumulative distribution function, we observed a striking resemblance between the network utilized in this study and the network model characterized by a power-law degree distribution. This finding aligns with the observations made by Albert et al. [19], who noted that power-law distributions are commonly observed in various real networks, including those describing intricate biological systems like the network employed in our analysis.

**Fig. 7:**
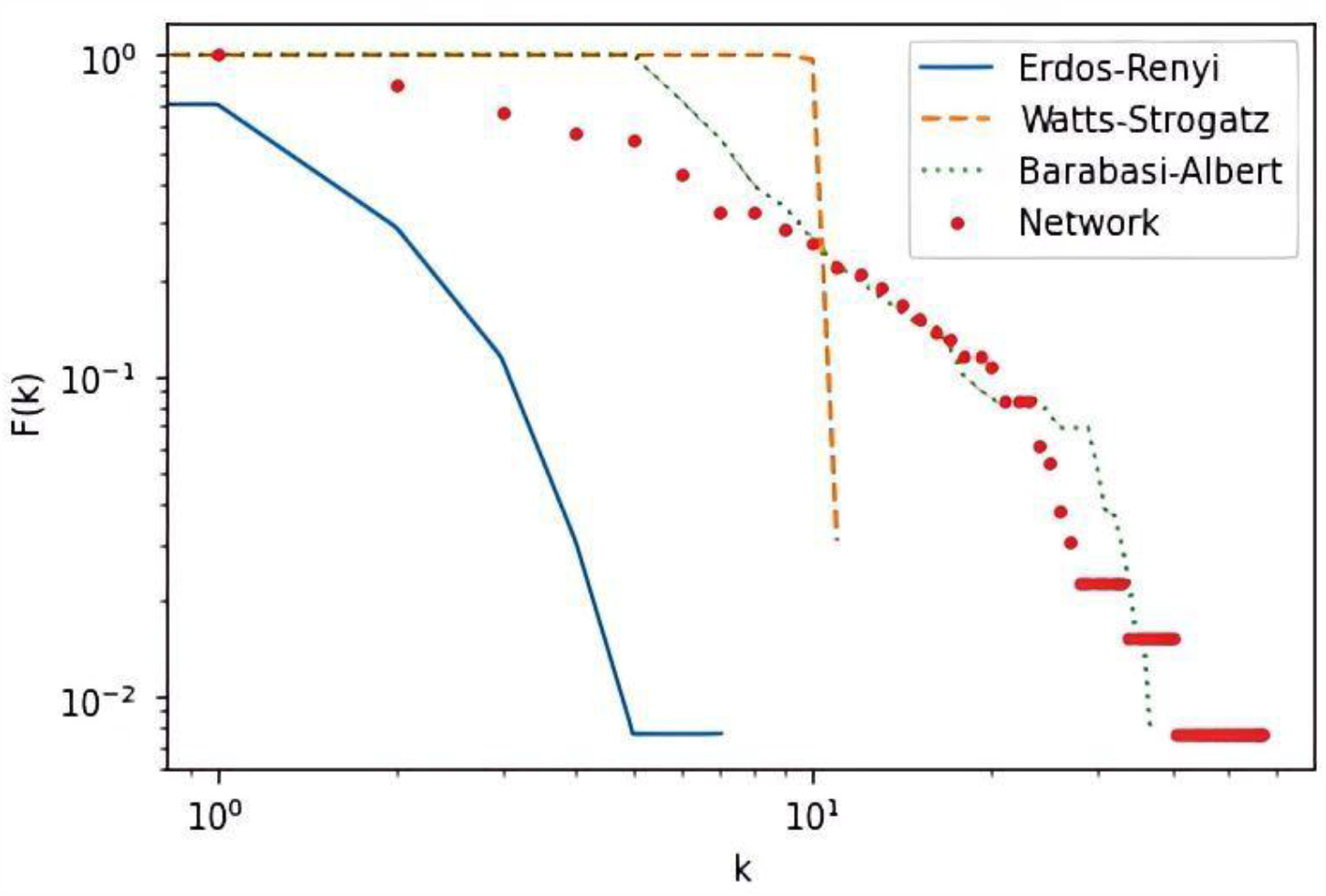
Convergence of our model with three canonical networks. Log-log plot of Erdos and Renvi (blue line), Watts and Strogatz (orange line), and scale-free networks (green line) networks compared to the experimental network of this study (red).

### 3 ATTRACTOR ANALYSIS

Upon examining the gene regulation network depicted in Figure 3, specifically in its unaltered state without any vertex inhibition, we observed that the percentage of genes satisfying the requisite conditions for transitioning from the malignant state to apoptosis was 29.6% for *body* samples and 14.8% for *protrusion* samples. This finding indicates an unfavorable configuration for the cell apoptosis process, as only two genes from the Bcl-2 family demonstrated a pro-apoptosis role. Moreover, the fact that crucial genes such as *CASP3, CASP6, and CASP7*, which play a fundamental role in the intrinsic apoptosis pathway, were inactive further supported the unsuitability of the configuration. Additionally, *XIAP* and *DIABLO*, which serve as inhibitors of *CASP9* and the IAP family, failed to meet the necessary conditions for initiating an apoptosis process, as illustrated in Figure 8.

**Fig. 8:**
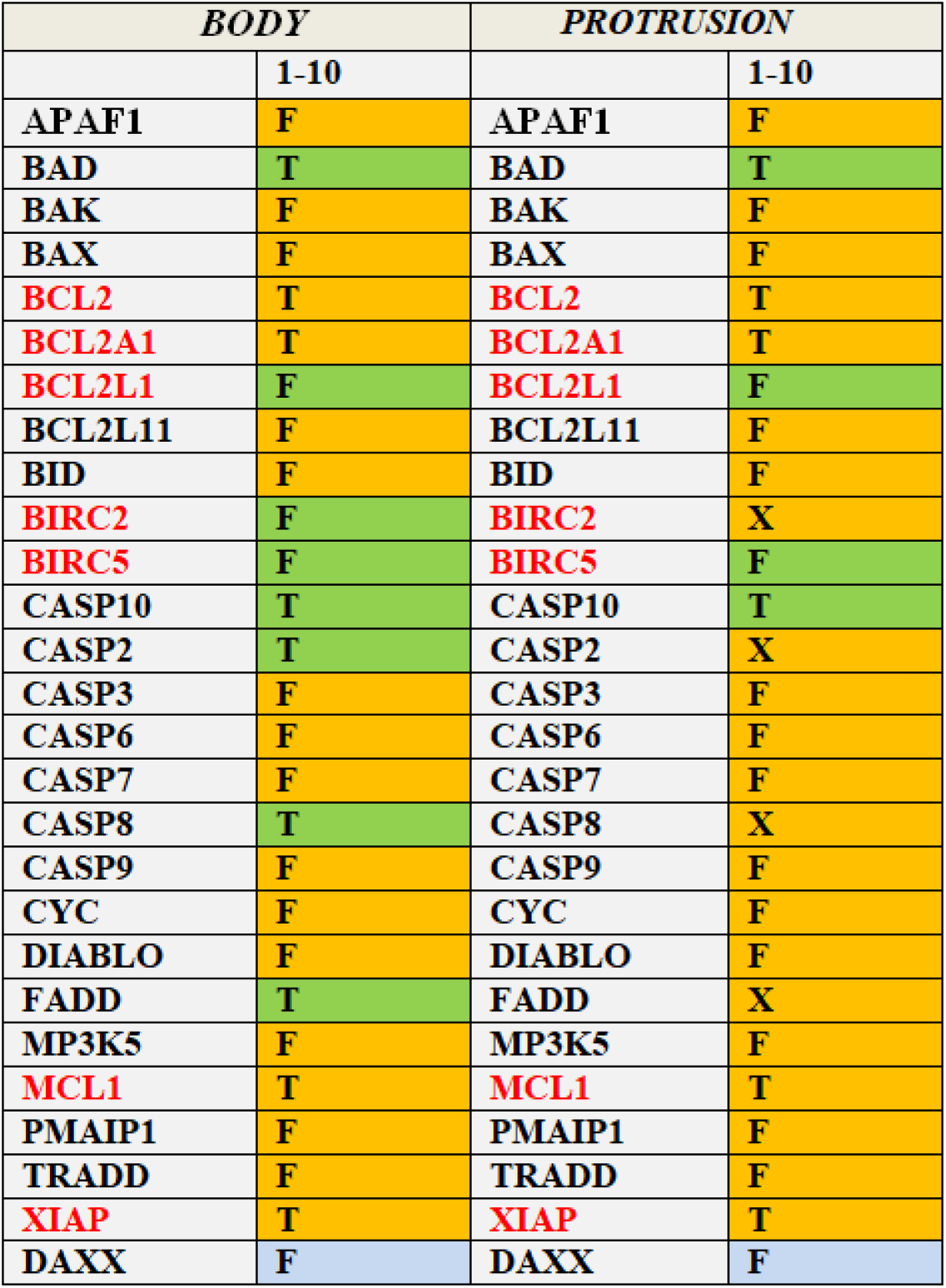
Configuration of the apoptosis-related genes in the attractors of the twenty comparisons of two MDA-MB-231 sample types, *body*, and *protrusion*. The seven genes indicated in red favor a pro-apoptotic mechanism if the gene is inhibited. On the other hand, the 20 genes displayed in black favor apoptosis if genes are activated (**Supplementary Table 1**). The *DAXX* gene is an exception, for which we did not find any characterization of its pro-apoptosis state in the literature. T (for True or activation), F (for False or inhibition), and X (for a continuous alternation between T and F) represent the boolean value of the 27 apoptosis-related genes within the detected attractors. T, F, and X background colors indicate if there are matches between the detected state and the ideal one of the corresponding genes for apoptosis induction. The green background means correspondence, while the orange one indicates a divergence.

To replicate the outcomes described in Tilli et al. [11], we conducted *in silico* the inhibition of the five bench targets known to induce cell death in the MDA-MB-231 cell line *in vitro*. In the *body* samples, the percentages of genes in the apoptosis attractors that assume the state of inhibition or activation supporting cell death were 74.1% and 55.6% in the comparisons 1 to 6 and 7 to 10, respectively. As for the *protrusion* samples, we found that 41% of the genes in the apoptosis phenotype state were observed in comparisons 1 to 6, while in comparisons 7 to 10, the percentage rose to 74%. These results are depicted in Figure 9.

**Fig. 9:**
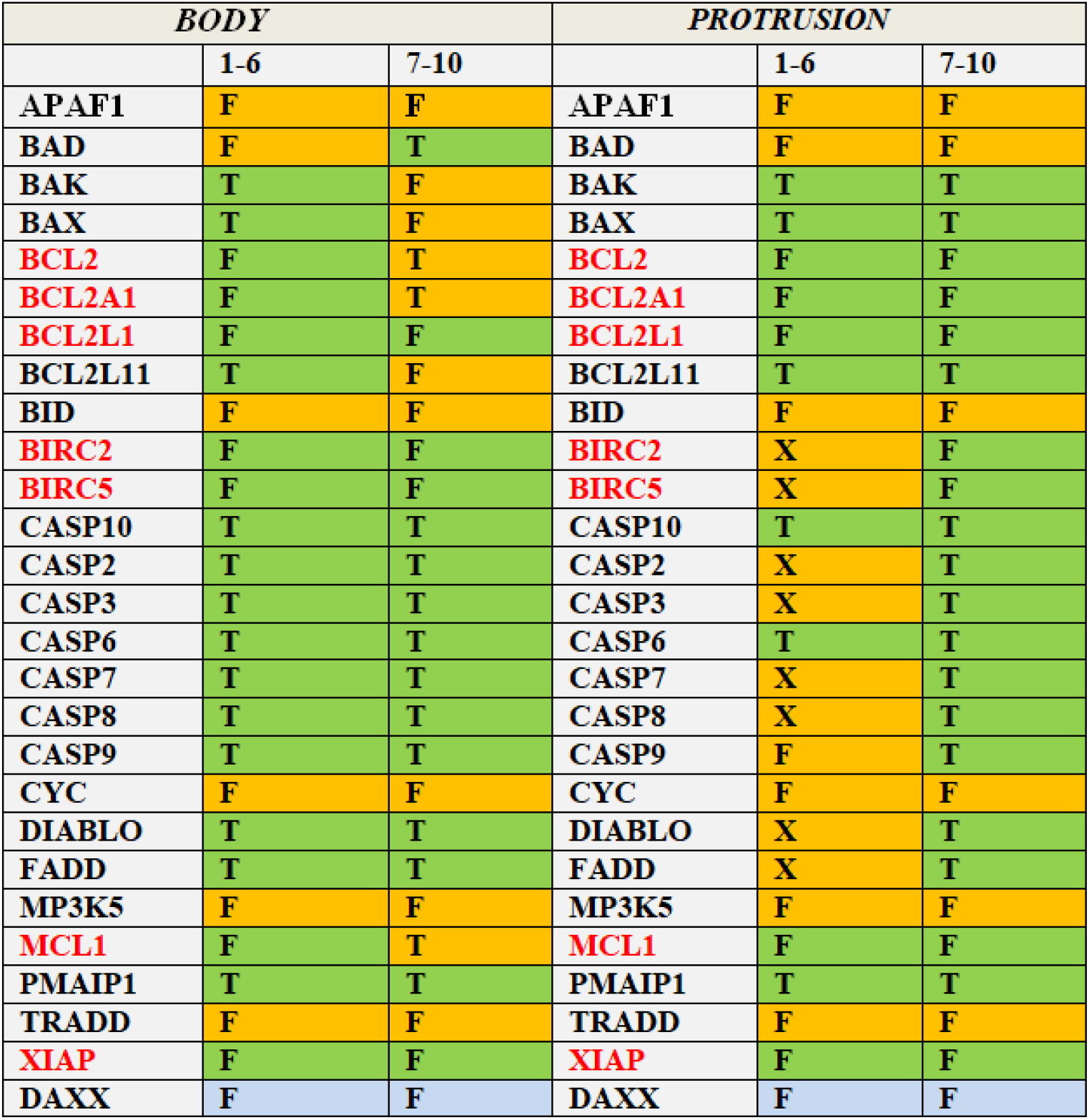
Attractors obtained by inhibiting the five bench targets: *CSNK2B, HSP90AB1, TK1, VIM*, and *YWHAB*. Columns *BODY* 1-6 and 7-10 refer to the ten *body* samples, while columns *PROTRUSION* 1-6 and 7-10 refer to the ten *protrusion* ones.

Comparing these results with those presented in Figure 8 (obtained without any gene inactivation in the network), there is a noticeable difference in both quantitative and qualitative aspects. Quantitatively, the percentages of genes aligned with the apoptosis state in the attractors are significantly higher, indicating a certain degree of representation of the *in vitro* experiment within the model. Qualitatively, the presence of the apoptotic state in the Bcl-2 and Caspase families and DIABLO and XIAP supports the expected outcomes of the bench experiment in the *body* 1-6 and *protrusion* 7-10 comparisons. However, in the *body* 7-10 and *protrusion* 1-6 comparisons, this alignment is only partially observed due to discrepancies in the RNA-seq profiles of the MCF10A cell lines used in these particular comparisons.

### 4 NETWORK MODULARITY ANALYSIS

The network modularity analysis revealed that the apoptosis-related genes tended to form distinct clusters with clear functional characteristics. This observation is depicted in Figure 10, where the two different groups of apoptosis-related genes are illustrated.

**Fig. 10:**
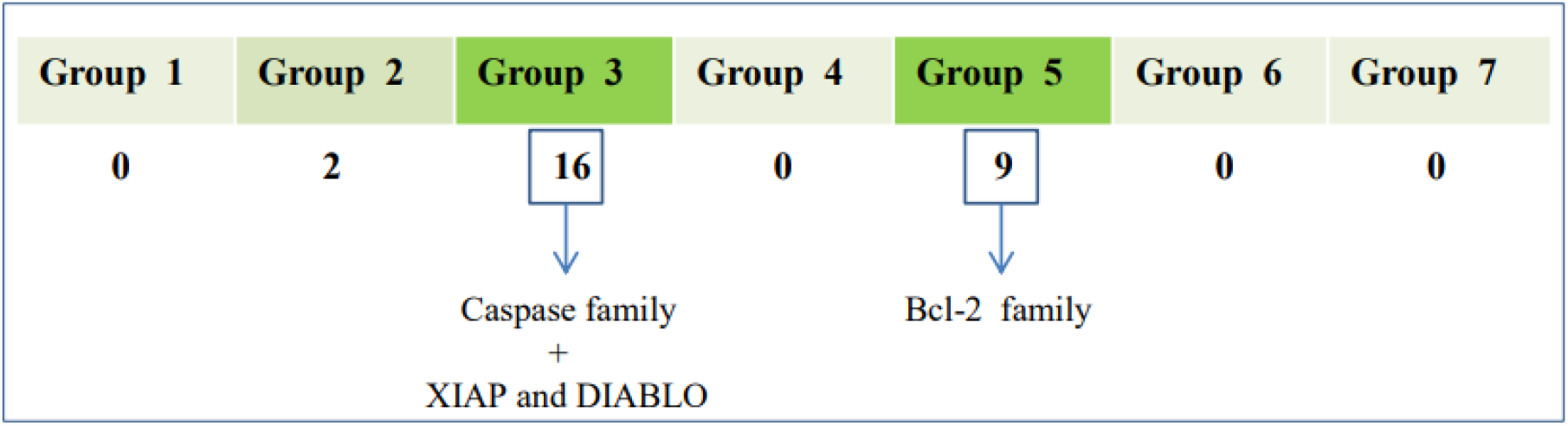
Network modularization process through the Clauset-Newman-More algorithm with the number of components of the apoptosis group in each of the seven modules. Groups 3 and 5, highlighted in green, indicate the groups belonging to the same modules relevant to the cell apoptosis process (mainly Caspases and Bcl-2 families) [29].

The majority of apoptosis-related genes, approximately 92.5%, are found in Groups 3 and 5. Group 3 comprises the complete *Caspases, XIAP*, and *DIABLO* group, while Group 5 includes the entire Bcl-2 family. The modular distribution of these genes, as depicted in **Supplementary Table S4**, demonstrates their tendency to cluster together. This characteristic is of great significance in the methodology employed in this study as it enables the utilization of a relatively small number of target vertices to activate these genes.

### 5 SHORTEST PATH EVALUATION

By examining the shortest path connecting the five bench targets with the apoptosis-related genes, we identified three genes present in at least one of these paths, namely HIF1A, XIAP, and BCL2. Since *XIAP* and *BCL2* are part of the apoptosis group, which serves as an indicator of the network’s state, and they also serve as the final nodes in the shortest paths, we did not evaluate the effects of inhibiting these genes on the apoptosis attractor. However, since *HIF1A* is not a member of the apoptosis group, we replaced *XIAP* and *BCL2* with their respective input nodes, *STAT5A* and *BRCA1* (Figure 11).

**Fig. 11:**
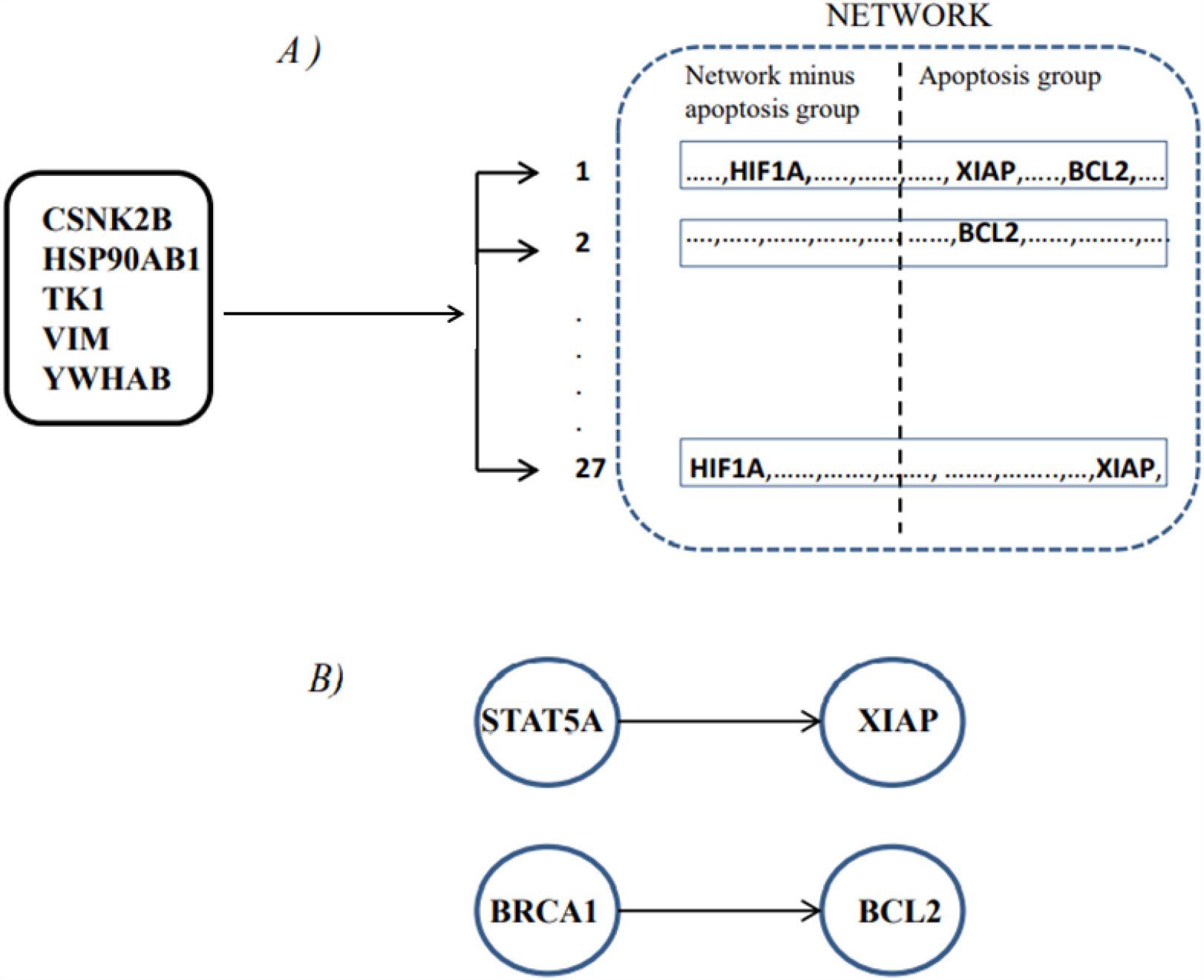
Process of new target identification by the shortest search. **Panel A:** *HIF1A, XIAP*, and *BCL2* are present in at least one of the 27 shortest paths. From these three genes, only *HIF1A* did not belong to the apoptosis-related group. **Panel B:** *STA5A* and *BRCA1* represent the only input genes of *XIAP* and *BCL2*. Thus, *HIF1A, STAT5A*, and *BRCA1* were the optimized target nodes detected within the shortest paths between the five bench targets and the 27 apoptosis-related genes. These three vertices were excellent candidates to complete the new set of optimized target genes that trigger the network within the apoptosis state.

### 6 OPTIMIZING THE NUMBER OF TARGETS

Identifying the vertices with the highest centrality between the five bench targets and the 27 apoptosis-related genes allowed us to explore the impact of inhibiting the new targets on the apoptosis attractor and propose a novel approach for selecting therapeutic targets. By applying the algorithm outlined in Figure 5, we identified *STAT5A, BRCA1*, and *HIF1A* as highly central vertices. Hence, we targeted the inhibition towards these three genes instead of inhibiting CSNK2B, HSP90AB1, TK1, VIM, and YWHAB. Combining these three genes successfully activated the apoptosis attractor in all sample comparisons. The outcomes achieved by inhibiting *HIF1A, STAT5A*, and *BRCA1* as substitutes for the five bench targets are given in Figure 12.

**Fig. 12:**
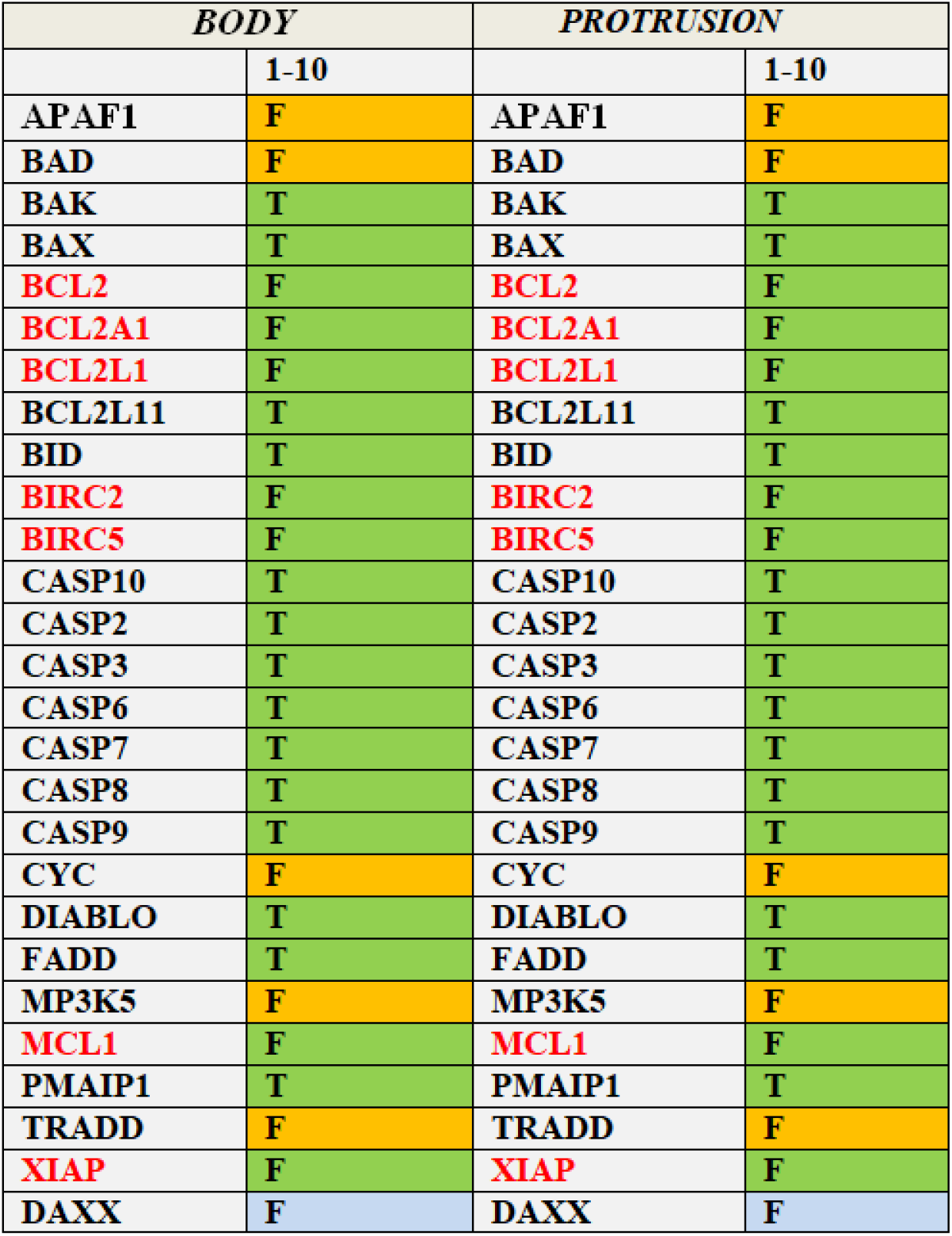
Activation or inhibition status detected in the 27 genes constituting the apoptosis-related group in the attractors of 10 *body* (1-10) and ten *protrusion* (1-10) sample comparisons by inhibiting *HIF1A, STAT5A*, and *BRCA1* instead of *CSNK2B, HSP90AB1, TK1, VIM*, and *YWHAB*.

In Figure 12, we demonstrated the simultaneous induction of cell apoptosis in both *body* and *protrusion* types for the *Bcl-2* and *Caspase* gene families across all combinations of RNA-seq data. Notably, *BID*, a pro-apoptosis member of the *Bcl-2* family, is activated in this simulation. The activation of BID plays a crucial role in activating downstream Caspases by directly activating *BAX* and *BAK*. This activation is absent in simulating the apoptosis attractor using the five bench genes (Figure 8). Similar considerations apply to *DIABLO*, which exhibited inconsistent expression in the six RNA-seq protrusion combinations from 1 to 6 (Figure 8).

Consequently, we simulated the induction of the network into the apoptosis attractor using various gene inhibition strategies. By inhibiting the five bench targets, we achieved a configuration conducive to apoptosis in a significant portion of the genes within this group (shown in green on the graph) (Figure 13). The percentage of genes activated in the apoptosis-related group was notably higher (74.1%) compared to scenarios without bench target inhibition (first column). When *HIF1A, STAT5A*, and *BRCA1* were selected as targets for inhibition, the proportion of genes activated in the apoptosis group further increased (77.8%) and remained consistent regardless of the MDA-MB-231 fraction or the MCF10A RNA-seq data used to identify the up-regulated genes in MDA-MB-231 (Figure 13, right column).

**Fig. 13:**
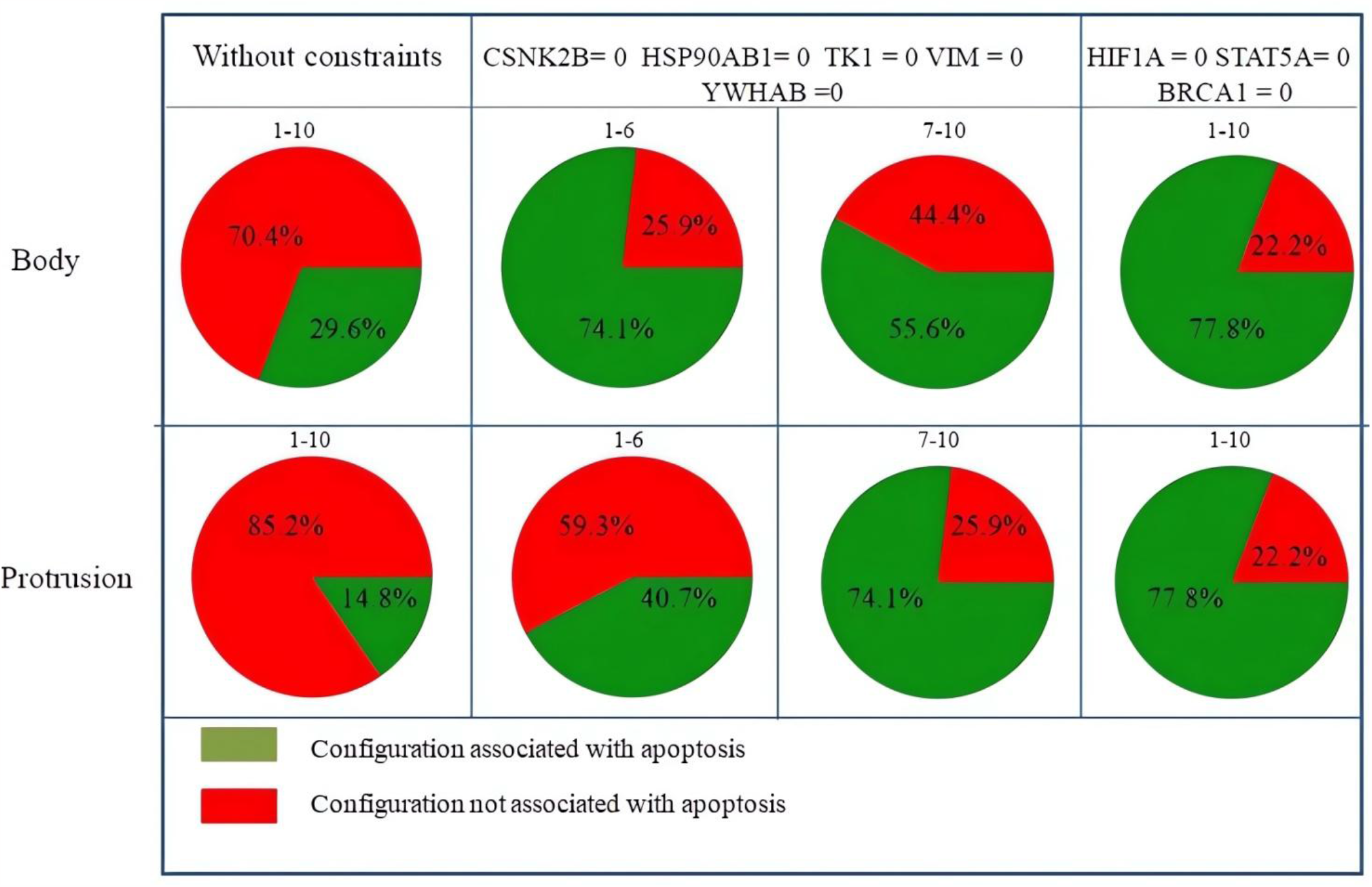
The results detected on the *body* and *protrusion* fractions of MDA-MB-231 used in this report, respectively. The columns identify which network genes were kept silenced in the dynamic simulation of the model. The green color, as opposed to the red one, indicates the percentage of the genes of the apoptosis group presenting a stable configuration of activation or inhibition associated with the attractor of the apoptosis phenotype.

Based on the findings above, it might be inferred that the transition from sample-specific malignant basins of attraction in *body* and *protrusion* samples, respectively, occurred towards a unified basin of attraction that signified a generalized state of cellular apoptosis, which was achieved by inhibiting *HIF1A, STAT5A*, and *BRCA1* (Figure 14).

**Fig. 14:**
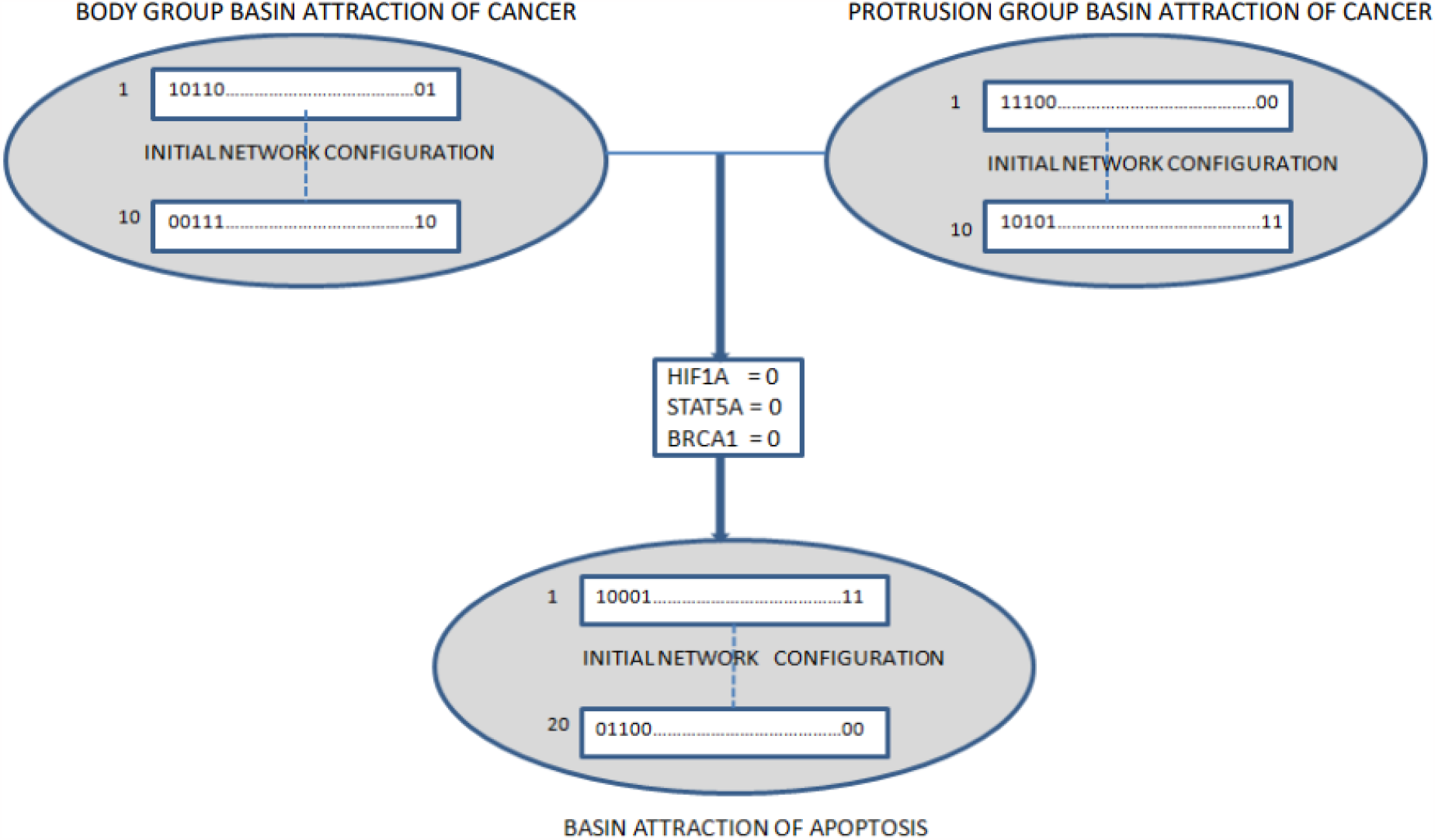
Boolean description of the transition from two basins of attraction representing the malignant cellular state of *body* and *protrusion* to a single basin of attraction of the cellular apoptosis state, obtained by inhibiting *HIF1A, STAT5A*, and *BRCA1*.

## DISCUSSION

As a multifaceted disease, cancer is influenced by numerous factors that cannot be comprehensively understood solely through molecular analysis. Consequently, there is a growing inclination to integrate molecular data with the dynamic characteristics of biological networks, employing computational and mathematical modeling techniques to gain deeper insights into the underlying biological mechanisms driving its progression [30].

The choice between quantitative and qualitative modeling approaches depends on the nature of the available data. Quantitative modeling, which involves ordinary differential equations and requires kinetic parameters, becomes challenging and feasible only for gene regulatory networks of limited scale [31]. In contrast, qualitative Boolean network modeling provides a viable alternative, allowing for relatively straightforward dynamic simulation of complex biological systems [32]. This approach proves beneficial in exploring regulatory interactions in protein expression [33] and developing strategies for therapeutic interventions [34]. It is also important to note that since the model proposed in this work is a Boolean-type model, we implicitly assume that the values of the system components are binary and Boolean functions govern their interaction. Such a description of the system dynamics, termed qualitative, necessarily implies a loss of the functional detail of the system that a quantitative methodology can provide instead. In addition, having chosen a synchronous rather than asynchronous network update system, giving preference to the deterministic nature of interactions an easy interpretation of results, a rough approximation was accepted in the timing mechanisms of the system elements, at the expense of the stochastic nature of these interactions.

In addition to the documented characteristics, we conducted functional compatibility checks to validate the Boolean model used in this study against the results obtained from an *in vitro* experiment [11] involving silencing five genes using siRNA. This experiment induced apoptosis in the MDA-MB-231 cell line. Our data show that our system can generate functionally compatible outcomes by inhibiting the same genes as in the *in vitro* experiment. Thus, we successfully replicated the behavior of an actual biological system within the Boolean dynamics of the gene regulatory network implemented in our research.

In this study, we utilized a Boolean network that represents a set of up-regulated genes in breast cancer to identify attractors corresponding to specific cellular phenotypes. We further assessed the compatibility of the Boolean network with an existing biological system by comparing it with an *in vitro* experiment [11]. The assignment of Boolean values to the network nodes followed the algorithm depicted in Figure 2. This algorithm facilitated the Booleanization of RNA-seq values based on the gene expression variations between malignant and non-malignant cell lines. By considering the gene expression differences across different cell lines, we were able to perform dynamic simulations of our model on a substantial number of comparisons, yielding valuable insights. Indeed, rather than solely relying on a single control for the four malignant samples (two *body* and two *protrusion*) to Booleanize the RNA-seq values and identify attractors, we expanded our approach by incorporating five samples of the non-malignant cell line MCF10A. By incorporating these additional samples, we derived gene expression differences that enabled us to perform a more comprehensive analysis. This strategy resulted in twenty combinatorial comparisons, significantly enhancing the numerical significance of network configurations and enabling the application of the procedures outlined in this study.

Considering cancer phenotype as basins of attraction in the epigenetic landscape [35], this report aimed to cause the transition from a basin of attraction of malignant type to that of apoptosis [36] through the perturbation of a subset of genes belonging to the network. For this purpose, we developed an algorithm (Fig. 5) that optimizes the choice of the network elements to produce a transition from one specific phenotype to another.

The approach of exploring the relationship existing between the results of an *in vitro* experiment, the insertion of a specific group of genes for apoptosis into the system, and the investigation of the network structure through the analysis of shortest paths between the five bench targets and the apoptosis-related group, represents an innovative strategy for clinical applications to increase patient benefit in personalized approaches of cancer therapies.

Including the apoptosis-related gene group within the network was a reference to evaluate the induction of cell death attractors through vertex inhibition. The results obtained in this study, depicted in Figure 13, schematically illustrate the effectiveness of this methodology. By inhibiting the three genes *HIF1A, STAT5A*, and *BRCA1*, we observed a transition in the system dynamics from malignant basins of attraction to those associated with cell apoptosis across all analyzed samples. The comparison of this result (Figure 12) with that obtained by reproducing the *in vitro* experiment (Figure 9), shows the robustness of the data obtained by applying the procedure of Figure 5. Inhibiting the five genes (*CSNK2B, HSP90AB1, TK1, VIM, YWHAB*) described in Tilli et al. [11] promotes a configuration of the 23 apoptosis-related genes conducive to apoptosis. However, the quantitative uniformity of this configuration varies among different comparisons. In the *body* sample, lines 1 to 6 exhibited a greater inclination towards apoptosis, whereas lines 7 to 10 in the *protrusion* sample displayed a higher favorability towards apoptosis (Figure 9). Conversely, when inhibiting the three genes (*HIF1A, STAT5A, BRCA1*) as detected through the procedure outlined in

Figure 5, a more consistent profile of activated apoptosis-related genes was observed across all comparisons (Figure 12). Despite the divergent attractors between the body and protrusion samples, the induction of apoptosis by inhibiting *HIF1A, STAT5A*, and *BRCA1* underscores the method’s robustness.

According to Figure 11, the target genes identified through the procedure outlined in Figure 5 should ideally be *HIF1A, XIAP*, and *BCL2*, considering our objective of identifying target genes capable of activating or inhibiting the 27 apoptosis-related genes, regardless of their specific configuration. Consequently, we decided to avoid designating *XIAP* and *BCL2* as targets. One possible alternative was substituting them with their respective input nodes, *STAT5A* and *BRCA1*, which do not belong to the apoptosis group. Therefore, it cannot be ruled out that the combined inhibitory effect on the *HIF1A, XIAP*, and *BCL2* genes may significantly induce apoptosis in cancer cells.

The role of *HIF1A, STAT5*, and *BRCA1* is well documented in tumors. *HIF1A* encodes the HIF-1α protein, whose level is regulated by hypoxia and other mechanisms, and is part of the heterodimeric transcription factor HIF-1. HIF-1α has crucial roles in many tumorigenic processes, such as epithelial-mesenchymal transition (EMT), metastasis, cancer cell metabolism, and angiogenesis [44, 45, 46]. Interestingly, there is a crosstalk between the HIF-1 and p53 pathways to determine cell fate depending on hypoxic conditions [47, 48]. Therefore, targeting the HIF-1 signaling in cancer can be a promising therapeutic strategy [46].

The transcriptional factor STAT5 is a member of the JAK-STAT (Janus kinase/Signal transducer and activator of transcription) pathway, which is altered in many tumors. Activated STAT5 upregulates the expression of genes involved in cell proliferation, invasion, angiogenesis, and the inhibition of apoptosis [49]. The exact role of STAT5 in breast cancer is still under debate. The STAT5 activation in tumor macrophages by derived factors from breast cancer cells led to the expression of anti-tumor immune stimulatory genes [50]. On the other hand, it was shown that STAT5a, an isoform of STAT5, could confer resistance to doxorubicin [51] and combined PI3K/mTOR and JAK2/STAT5 pathways inhibition induced cell death in triple-negative breast cancer [52].

*BRCA1* is an essential gene in DNA repair and cell cycle regulation. When mutated, the risk of developing many cancers significantly increases, especially for breast and ovarian tumors [53]. Several studies have shown increased brain metastasis frequency in patients carrying *BRCA1* mutations [54, 55]. Another fundamental role of this gene is the maintenance of genomic stability [56]. Therefore, *BRCA1* is essential to tissue homeostasis.

More specifically, several reports have established the relationship between the inhibition of *HIF1A, STAT5A*, and *BRCA1* genes and the induction of apoptosis in the MDA-MB-231 cell line. In the case of *HIF1A*, suppressing its expression using siRNA has been shown to inhibit cell growth and enhance apoptosis [37]. Inhibition of *STAT5A* has been correlated with reduced metastasis and growth of breast cancer tumor cells [38]. Additionally, the knockdown of *STAT5A* restores cellular sensitivity to TRAIL-induced apoptosis [39]. As for *BRCA1*, its RNAi-mediated silencing, along with miR-342 transfection, has been found to increase the percentage of apoptotic cells [40]. Furthermore, *BRCA1*-depleted MDA-MD-231 cells exhibited heightened susceptibility to proteasome inhibitors [42]. Considering the known functions and the consequences of the deregulation of these three genes in cell homeostasis, our study underscores the impact of inhibiting them on promoting apoptosis induction in the MDA-MB-231 cell line. It is important to note the interplay between HIF-1 and p53 pathways to determine cell death under hypoxic conditions [47, 48]. The inhibition of HIF1A could favor p53 in its apoptotic roles. However, in our model, p53 was inactivated since it is mutated in this cell line and not working as a tumor suppressor (41). Therefore, other mechanisms need to be investigated more deeply in the future.

The outcomes presented in this study hinged on the fine-tuning of the transfer functions (eqs. 6-10) to align the model with the *in vitro* experiment [11]. However, the concurrence observed between the *in vitro* results, and the computational simulation indicated a satisfactory level of model representativeness, warranting its potential for future optimization and application in therapeutic scenarios. Consequently, it becomes feasible to integrate specific experimental findings with computational hypotheses formulated to tackle therapeutic challenges associated with cancer.

It is worth noting that the identified therapeutic targets are the results obtained by running the algorithm presented in Figure 5 on the boolean network model constructed and validated by our group. The results obtained through their inhibition show that their choice is necessary and sufficient to achieve optimization in qualitative terms of the performance obtained from the in vitro experiment taken as a reference in this report. All this does not exclude the possibility of not having considered other important therapeutic targets that have emerged in other contexts.

The main objective of our research was to identify therapeutic targets on which an inhibition action is capable of causing a change in the state trajectory of the system, consequently producing a change in the system’s final target attractor.

However, the use of Boolean gene regulatory networks in some research areas, such as pharmacogenetics [43], can be challenging in identifying the complicated mechanisms between the genome, its products (RNAs and proteins), and the cellular-level response to drugs.

Because of the complex interactions that exist among molecules involved in a carcinogenic process, a perturbation analysis method such as the one we used in our research can be useful in dealing with such complexity, proposing specific interventions on the system by guiding and facilitating the subsequent choice of therapy useful for the purpose. Indeed, once therapeutic targets have been identified, there is the possibility of pharmacologically acting on them directly or through signaling pathways in which they are involved, through drugs currently in use.

Another therapeutic possibility of greater complexity is using siRNA molecole encapsulated in nanoparticles specific to the identified targets.

The outcomes presented in this study are derived from the analysis of data obtained from specific biological samples. The growing abundance of such information on distinct pathological conditions of cancer highlights the versatility of our model in accommodating various configurations of the same disease. The positioning of the method developed in this study within personalized medicine reflects its capacity to address individualized approaches to cancer treatment.

## CONCLUSION

In this research, we implemented a new computational method for optimizing the number of potential targets for breast cancer. We constructed a Boolean Gene Regulatory Network Model of a breast cancer tumor and validated it using RNA-seq data from tumoral cell lines. We achieved these results by integrating experimental data with those obtained from an extensive literature search in Boolean gene regulatory networks, for which the analysis of the corresponding attractors allowed the identification of potential therapeutic targets. In future work, we intend to apply our method to actual patient data to validate our results in the context of personalized medicine.

## DATA AVAILABILITY STATEMENT

Source code and supplementary data used in this study are available at: https://github.com/Domenico321/therapeutic-optimization

